# MicroRNA-146a Deficiency Protects NOD Mice from Autoimmune Diabetes by Enhancing c-Rel-Dependent Regulatory T Cell Function

**DOI:** 10.64898/2026.09.23.753219

**Authors:** Corynn N Appolonia, Shrikanth Basavarajappa, Angela Rose Liu, Katie Northrop, Parameswaran Ramakrishnan

**Affiliations:** Department of Pathology, Case Western Reserve University, 6526 Wolstein Research Building, 2103 Cornell Road, Cleveland, OH, 44106, USA; The Case Comprehensive Cancer Center, Case Western Reserve University, 2103 Cornell Road, Cleveland, OH, 44106, USA; Department of Biochemistry, Case Western Reserve University, 2109 Adelbert Road, Cleveland, OH, 44106, USA; University Hospitals-Cleveland Medical Center, 11100 Euclid Ave, Cleveland, OH, 44106, USA; Louis Stokes Veterans Affairs Medical Center, 10701 East Blvd, Cleveland, OH, 44106, USA

## Abstract

Type 1 diabetes (T1D) is a chronic autoimmune disease characterized by T-cell mediated destruction of pancreatic islet β-cells with genetic, environmental, and molecular triggers involved in disease pathogenesis. Patients with T1D have elevated serum levels of microRNA146a (miR146a). Polymorphisms in the miR146a gene that result in reduced expression of miR146a are associated with protection from T1D. We studied physiological regulators of miR146a expression and found that both hyperglycemia and elevated O-GlcNAcylation increased miR146a expression in T cells. Peripheral blood mononuclear cells (PBMCs) from T1D patients showed increased miR146a and O-GlcNAc transferase (OGT) expression, suggesting increased O-GlcNAcylation may promote miR146a expression in T1D patients. To determine the genetic and developmental role of miR146a in T1D, we generated miR146a-knockout (KO) non-obese diabetic (NOD) mice and found that absence of miR146a significantly protected NOD mice from spontaneous autoimmune diabetes. While we found no impact of miR146a knockout on general hematopoietic parameters and immune cell populations, remarkably, immune cell infiltration into the pancreas was significantly attenuated. Protection from autoimmune diabetes in miR146a-KO NOD mice was associated with increased regulatory T (Treg) cells in the spleen and pancreatic lymph node. Mechanistically, absence of miR146a increased NF-κB c-Rel expression in Treg cells and enhanced c-Rel binding at the Forkhead box protein P3 (FOXP3) promoter, which positively regulated Treg cell development and suppressor function, offering protection from T1D in miR146a-KO NOD mice. Our findings reveal miR146a as a key regulator of Treg cell-mediated immune tolerance through controlling NF-κB c-Rel-dependent FOXP3 expression and suggest targeting miR146a as a potential strategy to restore peripheral tolerance in T1D.

## Introduction

Type 1 diabetes (T1D) is a chronic autoimmune disease characterized by immune cell-mediated destruction of pancreatic islet β-cells, leading to insulin deficiency and hyperglycemia. The autoimmunity in T1D results from a loss of self-tolerance to pancreatic β-cell antigens. Regulatory T (Treg) cells play a major role in maintaining self-tolerance and preventing the initiation of autoimmune diseases (Rajendeeran and Tenbrock 2021). In patients with T1D, Treg cell frequency and function are altered, preventing maintenance of self-tolerance (Hull, Peakman, and Tree 2017). As a result, aberrantly activated autoreactive T cells from the pancreatic lymph node migrate into the pancreatic islets, resulting in β-cell destruction (Sandor, Jacobelli, and Friedman 2019; Turley et al. 2005).

The mechanisms behind autoimmunity in T1D are complex and multi-factorial. Recent studies have implicated various microRNAs in the pathogenesis of T1D. MicroRNAs are short, single-stranded RNA molecules that are involved in post-transcriptional regulation of gene expression. MicroRNAs classically interact with the 3’ untranslated region (UTR) of target messenger RNAs (mRNAs) based on complementarity. This interaction mostly commonly suppresses translation or promotes degradation of the target mRNA (Gebert and MacRae 2019). Dysregulated microRNA expression has been associated with pancreatic β-cell dysfunction, decreased insulin synthesis, and increased autoimmunity (Margaritis et al. 2021).

MicroRNA146a (miR146a) dysregulation has been implicated in both T1D patients and in mouse models of T1D. In the non-obese diabetic (NOD) mouse model of autoimmune diabetes, miR146a significantly increases in the islets of mice with age and peri-insulitis (Roggli et al. 2010). In humans, polymorphisms in the miR146a gene (rs2910164) are associated with decreased miR146a expression and protection from T1D (Assmann et al. 2017). Additionally, plasma levels of miR146a are significantly increased in T1D patients at diagnosis compared to sex and age matched controls (Garavelli et al. 2020).

miR146a is highly expressed in immune cells (Boldin et al. 2011). In T cells, miR146a is increased after T-cell receptor activation and in splenic T cells compared to thymic T cells (Yang et al. 2012). CD4^+^CD25^+^ Treg cells have an approximately 9-20-fold increase in miR146a expression compared to naïve T cells and CD4^+^CD25^-^ cells (Yang et al. 2012; L. F. Lu et al. 2010). Expression of miR146a is controlled by a distinct promoter region with binding sites for the NF-κB family of transcription factors (Taganov et al. 2006; Iacona and Lutz 2019). miR146a targets the mRNAs of interleukin-1 receptor-associated kinase 1 (IRAK1) and tumor necrosis factor receptor-associated factor 6 (TRAF6), which are adaptor proteins in toll-like receptor (TLR) and cytokine signaling (Taganov et al. 2006; Mortazavi-Jahromi, Aslani, and Mirshafiey 2020). More recently, NF-κB c-Rel has been identified as a direct target of miR46a in B cells (Cho et al. 2018).

In this study, we determined the role of miR146a in autoimmune diabetes. We used publicly available data for the transcriptomics analysis of miR146a expression and showed its increased levels in peripheral blood mononuclear cells (PBMCs) of T1D patients. PBMCs from T1D patients demonstrated increased expression of the enzymes that control O-GlcNAcylation, O-GlcNAc transferase (OGT) and O-GlcNAcase (OGA), with a disproportionate increase in OGT relative to OGA, suggesting the presence of hyper-O-GlcNAcylation. Hyperglycemia and enhanced O-GlcNAcylation in T cells resulted in elevated miR146a expression. To study the *in vivo* role of miR146a, we generated a miR146a-knockout (KO) NOD mouse model and discovered absence of miR146a protects from spontaneous autoimmune diabetes. miR146a deletion results in increased frequency and suppressor function of Treg cells, which mechanistically connects to increased NF-κB c-Rel binding to the promoter of FOXP3 in miR146a-KO Treg cells. Our results suggest a model where miR146a and c-Rel constitute a regulatory axis controlling Treg cells during T1D autoimmunity and set the stage for miR146a inhibition as a therapeutic approach to improve self-tolerance in patients with T1D.

## Materials and Methods

### Reagents and antibodies

The following reagents were used in this study: Thiamet-G (MedChemExpress), phorbol 12-myristate 13-acetate (Sigma), Ionomycin (Sigma), OSMI-1 (Sigma). Antibodies against the following proteins were used in this study: O-GlcNAc RL2 (sc-59624, Santa Cruz Biotechnology), β-Actin (C4, Santa Cruz Biotechnology), c-Rel (D3B8S, Cell Signaling Technology), p105/p50 (D4P4D, Cell Signaling Technology), p65 (C20, Santa Cruz Biotechnology), p65 (L8F6, Cell Signaling Technology), and vinculin (7F9, Santa Cruz Biotechnology). For flow cytometry, fluorescent antibodies against the following proteins were used: CD4 (GK1.5, BioLegend), CD8 (53-6.7, BioLegend), CD3 (145-2C11, Miltenyi Biotec), TCR γδ (GL3, Miltenyi Biotec), CD11b (M1/70, BioLegend), CD11c (N418, BioLegend), CD19 (6D5, Miltenyi Biotec), and FOXP3 (150D, BioLegend). Zombie NIR Fixable Viability Kit (BioLegend) was used for live-dead analysis.

### Cell culture

Jurkat cells were cultured in RPMI 1640 medium containing 10% fetal bovine serum (FBS), 100 U/mL penicillin/streptomycin, and 1% L-glutamine. Primary T cells and Treg cells were cultured in RPMI 1640 medium containing 10% FBS, 100 U/mL penicillin/streptomycin, 1% L-glutamine, 1% HEPES, 1% nonessential amino acids, 1% sodium pyruvate, and 0.05 mM β-mercaptoethanol. Cells were incubated at 37°C with 5% CO_2_.

### Animals

Dr. David Baltimore, California Institute of Technology kindly provided C57BL/6 miRNA146a knockout (KO) mice. NOD/ShiLtJ mice (strain #001976) were purchased from Jackson Laboratories and maintained in-house. All mice were housed in specific pathogen-free conditions. miRNA146a-KO NOD mice were generated by successive back crossing of miRNA146a-KO C57BL/6 mice onto the NOD mice background for 18 generations and then intercrossed to generate homozygous knockout mice at Case Western Reserve University animal facility. All animal experimental procedures were performed in accordance with the National Institutes of Health guidelines and regulations approved by the Institutional Animal Care and Use Committee of Case Western Reserve University under protocol #2013-0134. With the exception of diabetes development studies, all mice were 6 to 10 weeks of age at the time of experiments.

### PCR genotyping

The HotSHOT method (Truett et al. 2000) was used to isolate DNA from mouse ear snips. Genotypes were confirmed with PCR using DreamTaq PCR Master Mix (ThermoFisher) following manufacturer’s instructions and a three-primer pair system:

5’-ATCGCGGCCGCTTTAAGTGTAGAGAGGGGGTCAAGTA-3’;

5’-ATTGCTCAGCGGTGCTGTCCATCTGCACGA-3’;

5’-CTTGGACCAGCAGTCCTCTTGATGCACCTT-3’. PCR products were resolved on a 1% agarose gel and visualized with ethidium bromide staining.

### Blood glucose measurements

Diabetes development in the mice was monitored by measuring the blood glucose level following tail venipuncture using commercially available glucometer (OneTouch Ultra or Contour). Blood glucose levels in NOD mice were monitored weekly starting at 8 weeks of age until 35 weeks of age. Testing was performed twice a week when the blood glucose level rose above 250 mg/dL. Mice showing three consecutive blood glucose readings of greater than 250 mg/dL were declared diabetic and euthanized.

### Hematology

Blood was collected in tubes precoated with 0.5 M EDTA, and differential blood count analysis was carried out using HemaTrue hematology analyzer (Heska).

### Histology

The pancreases from wildtype NOD and miR146a-KO NOD mice were isolated and fixed in neutral buffered 10% formalin solution. Formalin fixed tissues were paraffin embedded, sectioned, and stained with hematoxylin and eosin by histology cores at CWRU and Fred Hutchinson Cancer Center, Seattle. Two research personnel blindly scored the stained slides independently for pancreatic islet infiltrates. Representative islets were photographed using a Motic or Leica DM IL LED microscope.

### Primary cell isolation

Single cell suspensions were obtained from the mouse thymus, spleen, and pancreatic lymph nodes through tissue homogenization using the rubber-capped plunger of a 3-cc syringe. Single cell suspensions were obtained from the mouse pancreas using 1-mL pipette trituration as previously described (Perides et al. 2011). Cells were passed through 40 µm cell strainer and centrifuged at 450 g. Red blood cells were lysed using RBC lysis buffer.

### Flow cytometry

For staining surface markers, cells were incubated with respective, fluorescent-labeled antibodies in FACS buffer (phosphate buffered saline, 0.5% bovine serum albumin, 0.1% sodium azide). For intracellular FOXP3 staining, cells were first stained with surface marker antibodies, followed by fixation and permeabilization using True Nuclear Transcription Factor Buffer Set (BioLegend) and intracellular staining with FOXP3 antibody. Cells were washed with FACS buffer and analyzed using Accuri or Cytoflex flow cytometer.

### Magnetic sorting

The MojoSort Mouse CD4^+^CD25^+^ Regulatory T cell Isolation Kit (BioLegend) was used to isolate CD4^+^CD25^-^ T cells and CD4^+^CD25^+^ T regulatory cells from the mouse spleen following manufacturer’s instructions. The purity and viability of isolated cells was confirmed with flow cytometry.

### T Proliferation assay

CD4^+^CD25^-^ T cells were stained with 1 µM CFSE (BioLegend) at 37°C for 15 minutes. CFSE stained cells were stimulated in a plate coated with 2 ug/mL anti-CD3 and anti-CD28 antibodies (Biolegend) for 72 hours, with or without CD4^+^CD25^+^ Treg cells. Proliferation was assessed by flow cytometric analysis of CFSE dilution using Cytoflex flow cytometer (BD Biosciences).

### Real-time quantitative PCR

For microRNA expression experiments, total RNA, including small RNAs, was isolated from cells using mirVana miRNA isolation kit (ThermoFisher). RNA was quantified using a NanoDrop spectrophotometer. 10 ng of total RNA was used for cDNA synthesis using the TaqMan MicroRNA Reverse Transcription Kit (Applied Biosystems). Quantitative real-time PCR was performed using TaqMan Fast Advanced Master Mix (ThermoFisher) according to manufacturer’s instructions and TaqMan MicroRNA Assay reagents for miR146a (000468), RNU48 (001006) as a small RNA housekeeping control for human cells, and snoRNA202 (001232) as a small RNA housekeeping control for mouse cells (ThermoFisher). Gene expression was quantified as relative expression compared to the housekeeping control gene.

### Western blotting

For total cell lysate, cells were lysed in Triton lysis buffer (1.0% Triton-X100, 20 mM HEPES [pH 7.6], 150 mM NaCl, 1 mM EDTA, and protease inhibitor cocktail) for 30 minutes on ice. Samples were centrifuged at 12,000 g for 10 minutes at 4°C to clear debris. For nuclear and cytoplasmic fractionation, cells were lysed on ice for 15 minutes in hypotonic buffer (10 mM HEPES [pH 7.6], 10 mM KCl, 0.1 mM EDTA, 0.1 mM EGTA, and protease inhibitor cocktail) to isolate the cytoplasmic fraction. NP-40 was then added at a final concentration of 0.625%. Samples were vortexed and then centrifuged at 12,000 g for 30 seconds. Supernatants were collected as the cytoplasmic lysate fraction. The remaining pellet was washed once in hypotonic buffer and then lysed on ice for 30 minutes in hypertonic buffer (20 mM HEPES [pH 7.6], 400 mM NaCl, 1 mM EDTA, 1 mM EGTA, and protease inhibitor cocktail). Samples were then centrifuged at 12,000 g for 10 minutes at 4°C, and supernatants were collected as the nuclear lysate fraction. Protein concentrations in lysates were quantified using the Pierce BCA Protein Assay Kit (ThermoFisher). Normalized volumes of lysate were either used for oligonucleotide pulldown assay or were resolved on a 9% SDS-PAGE gel. Proteins were transferred to a nitrocellulose membrane and probed with indicated antibodies. Enhanced chemiluminescence assay (GenDEPOT) was used for visualization.

### Oligonucleotide pulldown assay

For the oligonucleotide pulldown assay, NaCl concentration of the hypertonic buffer was normalized to 150 mM. Nuclear lysates with normalized protein concentrations were then incubated with 1 μg of biotinylated, annealed oligonucleotides corresponding to the c-Rel binding site in the FOXP3 promoter (De Jesus et al. 2021; Long et al. 2009) and 6 μg of sheared salmon sperm for 30 minutes at 4°C with rotation. Neutravidin beads were then added, and samples were incubated with rotation at 4°C for an additional 2 hours. Beads were washed with buffer at 150 mM. Beads were then heated at 95°C with 1X Laemmli sample buffer and centrifuged at 10,000 rpm for 10 minutes. Supernatants were analyzed via western blotting. The following primer pair was used for the biotinylated annealed oligonucleotides: Forward: Bio5′-TGC GGC TTC CAC GCC GTG GTT TTT CTT CTC GGT ATA AAA GCA AAG TT-3’, Reverse: 5′-AAC TTT GCT TTT ATA CCG AGA AGA AAA ACC ACG GCG TGG AAG CCG CA-3’.

### Statistical analysis

Statistical significance between two groups was analyzed using a student’s t-test. Statistical significance between three or more groups was analyzed using a one-way ANOVA with multiple comparisons. Values were expressed as mean ± SEM and p-value < 0.05 was considered significant.

## Results

### miR146a expression is increased in type 1 diabetes patients and is regulated by glycemic and O-GlcNAcylation status

Newly diagnosed type 1 diabetes (T1D) patients show upregulation of miR146a in the plasma compared to healthy controls (Garavelli et al. 2020). To assess whether miR146a is upregulated in immune cells of T1D patients, we analyzed the expression of miR146a in peripheral blood mononuclear cells (PBMCs) using a publicly available dataset on the Gene Expression Omnibus (GEO) database. Compared to healthy controls, we found PBMCs of newly diagnosed type 1 diabetes patients had a significant increase in miR146a expression (Fig. 1*A*).

**Figure 1.**
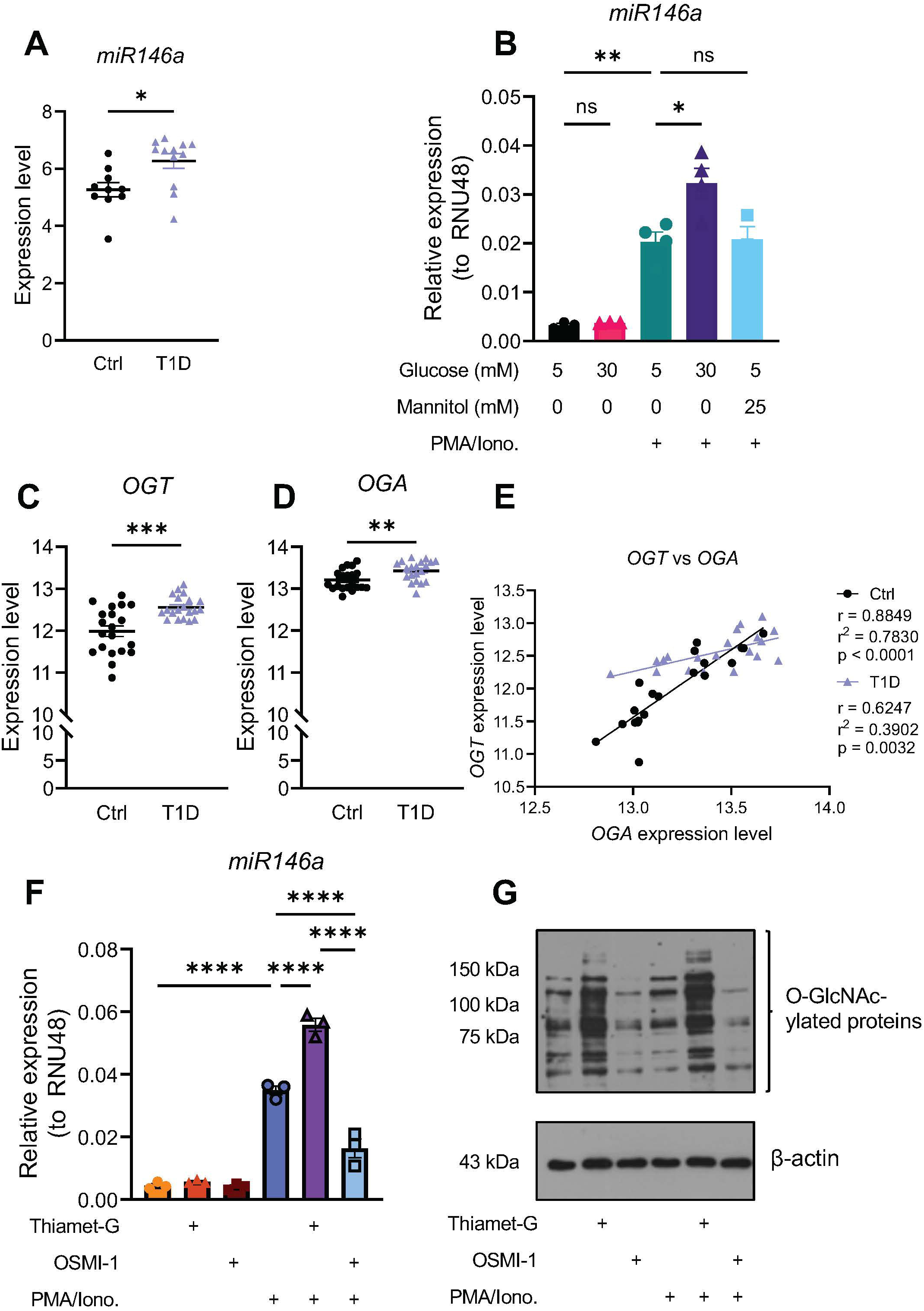
Elevated miR146a expression is observed in peripheral blood immune cells of T1D patients and in T cells exposed to hyperglycemia and enhanced O-GlcNAcylation. (A). MicroRNA146a expression in peripheral blood mononuclear cells (PBMCs) isolated from T1D patients and healthy controls was analyzed using publicly available microarray dataset GSE55099. Unpaired t-test, * p< 0.05. (B). Jurkat T cells were cultured for 48 hours in low-glucose media (5 mM glucose), high-glucose media (30 mM glucose), or mannitol osmotic control media (5 mM glucose + 25 mM mannitol). Cells were stimulated with 50 ng/mL PMA and 250 ng/mL Ionomycin or DMSO as a control for 8 hours. Samples were analyzed by RT-qPCR to determine the expression of miR146a relative to that of RNU48. One-way ANOVA with multiple comparisons. * p < 0.05, ** p < 0.01, ns = not significant. (C-D). MicroRNA146a expression in PBMCs isolated from T1D patients and healthy controls was analyzed using publicly available microarray dataset GSE193273. Unpaired t-test, ** p < 0.01, *** p < 0.001. (E). Scatterplot of OGT and OGA expression values for each healthy (black circle) and T1D (purple triangle) patient sample. r = Pearson correlation between OGA and OGT expression. Simple linear regression was performed for best-fit line. (F). Jurkat T cells were treated with 50 μM Thiamet-G or 25 μM OSMI for 24 hours. Cells were stimulated with 50 ng/mL PMA and 250 ng/mL Ionomycin or DMSO as a control for 8 hours. Samples were analyzed by RT-qPCR to determine the expression of miR146a relative to that of RNU48. One-way ANOVA with multiple comparisons. **** p < 0.0001. (G) Induction and inhibition of O-GlcNAcylation was assessed by western blotting using an antibody against O-GlcNAc (RL2). (A-D, F). Data is represented as mean ± SEM, and each data point represents a biological replicate.

miR146a expression is upregulated in T cells upon cellular activation in a NF-κB dependent manner (Yang et al. 2012). Other physiological regulators of miR146a expression in T cells that are relevant to T1D are unknown. Given increased expression of miR146a in T1D patients, we studied the effect of hyperglycemia on miR146a expression. Jurkat T cells were subjected to stimulation via phorbol 12-myristate 13-acetate (PMA) and ionomycin under physiological glucose (5 mM) and high glucose (30 mM) conditions. We observed significantly increased miR146a expression in cells exposed to hyperglycemia and T cell activation compared to T cell activation alone (Fig. 1*B*). The increase in miR146a expression was not due to high sugar-induced hyperosmotic stress as mannitol treatment did not result in an increase in miR146a expression (Fig. 1*B*).

One of the cellular effects of hyperglycemia is increased O-GlcNAcylation, a post-translational modification where a *N*-acetylglucosamine (GlcNAc) monosaccharide is added to serine or threonine residues on proteins (Mannino and Hart 2022). Glucose flux through the hexosamine biosynthesis pathway regulates the concentration of UDP-GlcNAc, which is the donor molecule for O-GlcNAcylation (Hart 2019). Thus, cellular O-GlcNAcylation levels are enhanced during hyperglycemic conditions (Hart 2019; Dennis et al. 2011; De Jesus et al. 2021). The addition and removal of O-GlcNAc is catalyzed by two proteins. The enzyme O-GlcNAc transferase (OGT) adds O-GlcNAc to proteins, while the enzyme O-GlcNAcase (OGA) removes O-GlcNAc from proteins (Mannino and Hart 2022). We found *OGT* expression was significantly upregulated in PBMCs from newly diagnosed T1D patients compared to healthy controls (Fig. 1*C*). We also found a modest increase in *OGA* expression in T1D PBMCs (Fig. 1*D*). We thus also assessed the relationship between *OGT* and *OGA* expression in healthy and T1D PBMCs using a correlation analysis. We observed a strong, positive correlation between *OGT* and *OGA* expression in healthy PBMCs (Fig. 1*E*), in accordance with previous studies that suggest OGA and OGT modulate the transcription of each other through cooperation with multiple transcription factors (Qian et al. 2018; Park et al. 2017; Kazemi et al. 2010). In contrast, in PBMCs from T1D patients, the correlation between *OGT* and *OGA* is weaker, with *OGT* expression consistently elevated despite variation in *OGA* expression. These findings suggest a loss of homeostatic regulation of *OGT* and *OGA* expression in T1D patients, resulting in a relative increase in *OGT* expression that may contribute to elevated protein O-GlcNAcylation in immune cells in T1D patients.

To investigate if O-GlcNAcylation regulates miR146a expression, we treated Jurkat T cells with PMA/Ionomycin and Thiamet-G, a chemical inhibitor of OGA, which will result in increased global cellular O-GlcNAcylation level. We observed a significant increase in miR146a expression in cells treated with Thiamet-G and PMA/Ionomycin (Fig. 1*F*). Next, we examined the effect of decreased O-GlcNAcylation by treating cells with OSMI-1, a chemical inhibitor of OGT. We found that OSMI-1 significantly reduces PMA/Ionomycin-induced miR146a expression (Fig. 1*F*). Reciprocal changes in total cellular O-GlcNAc levels after treatment with Thiamet-G and OSMI-1 were confirmed by western blot using an antibody against total O-GlcNAc (Fig. 1*G*). Taken together, these results suggest that hyperglycemia and enhanced cellular O-GlcNAcylation levels promote expression of miR146a in T cells, which could explain the observed increased expression of miR146a in PBMCs from type 1 diabetes patients.

### Generation of miR146a knockout mice on the NOD background

To study the role of miR146a in spontaneous autoimmune diabetes, we generated a miR146a knockout (KO) NOD mouse model (Fig. 2*A*). We crossed miR146a-KO mice on the C57BL/6 background with NOD wildtype (WT) mice to obtain miR146a^+/-^ heterozygous (H) mice. We repeatedly backcrossed heterozygous miR146 mice with NOD wildtype mice for at least 18 generations to obtain miR146a heterozygous mice on a congenic NOD background. This scheme included crosses of miR146 heterozygous females with wildtype NOD males as well to fix the Y chromosome. Finally, we mated miR146a heterozygous NOD mice to obtain miR146a^-/-^ knockout (miR146a-KO) NOD mice. Mouse genotypes were confirmed with genotyping PCR (Fig. 2*B*). Deletion of miR146a was further confirmed by determining miR146a expression via RT-qPCR (Fig. 2*C*). We analyzed the blood of WT and miR146a-KO NOD mice and did not observe any differences in major hematological parameters or peripheral blood immune cell percentages (Fig. 2*D*, Supplemental Table 1).

**Figure 2.**
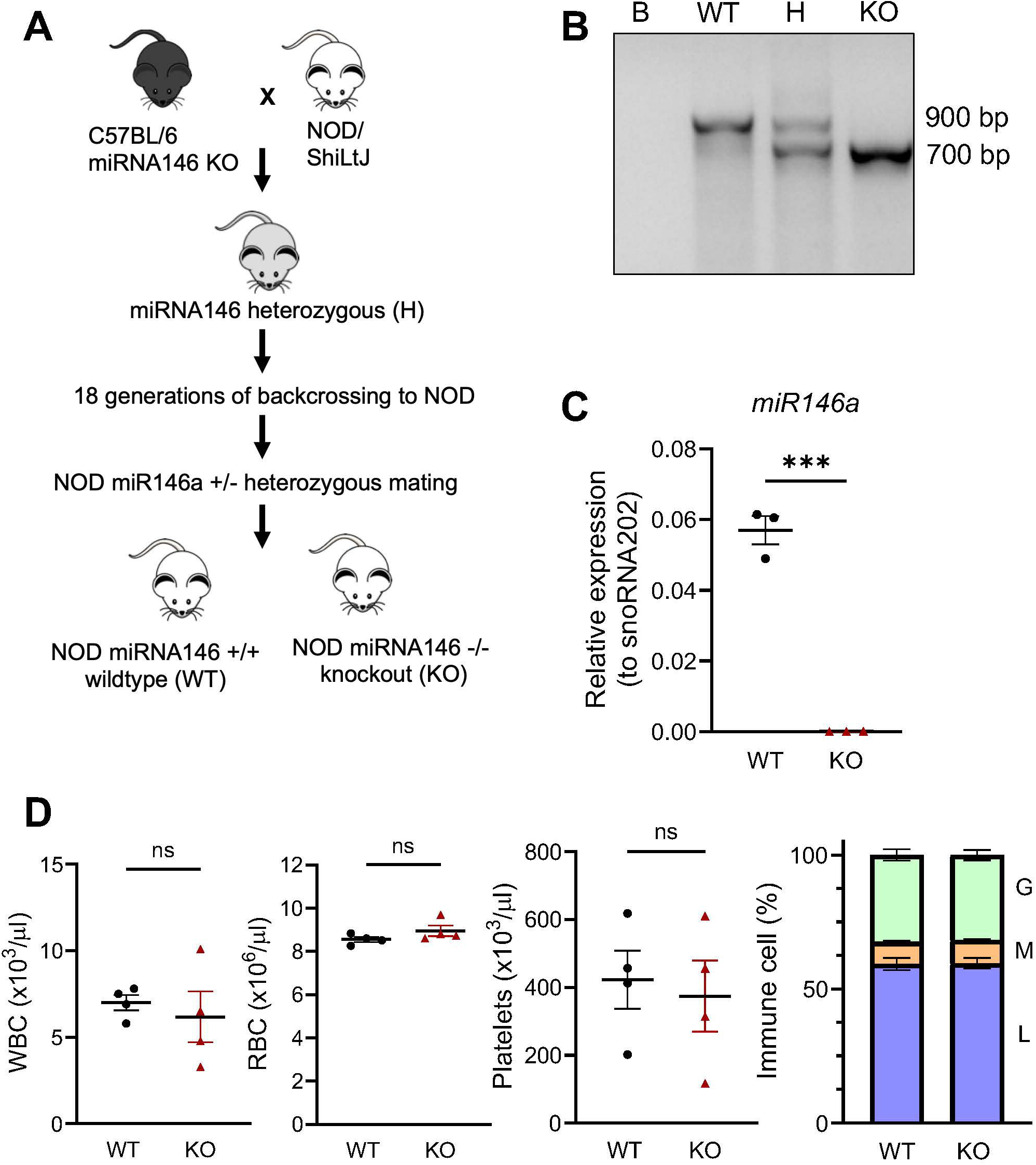
miR146a knockout on the NOD background does not affect general hematopoietic parameters. (A). Schematic demonstrating breeding strategy for generation of miR146a knockout NOD mice. (B). DNA was isolated from ear snips, and PCR was used to genotype wildtype (WT), miR146a heterozygous (H), and miR146a knockout (KO) NOD mice. Water (B) was used as a negative control. (C). CD4^+^ T cells were isolated from splenocytes of WT and miR146a-KO NOD mice, and small RNAs were isolated. Samples were analyzed by RT-qPCR to determine the expression of miR146a relative to that of snoRNA202. Unpaired t-test, *** p < 0.001. (D). Blood samples from wildtype (WT) NOD and miR146a knockout (KO) NOD mice were analyzed in the Hemavet blood analyzer. Immune cell percentages were calculated for granulocytes (G), monocytes (M), and lymphocytes (L). (C-D). Data is represented as mean ± SEM for n=3-4.

### miR146a deletion does not affect major immune cells in the thymus, spleen, and pancreatic lymph node in NOD mice but results in decreased T cells in the pancreas

Given the high expression of miR146a throughout immune cells compared to non-immune tissues (Boldin et al. 2011), we conducted immunophenotyping of the spleen and thymus in WT NOD and miR146a-KO NOD mice to further characterize our novel NOD mouse model. Analysis of age-matched 6–10-week-old WT and miR146a-KO NOD mice revealed no differences in the percentages of thymic CD4^+^CD8^+^ double positive lymphocytes and single positive lymphocytes (Fig. 3*A*). Similarly, we did not observe any significant differences in the percentages of splenic CD4^+^, CD8^+^, and γδ^+^ lymphocytes (Fig. 3*B*). Furthermore, no differences were observed in percentages of splenic CD19^+^ B cells, CD11b^+^ cells, and CD11c^+^ cells (Supplemental Fig. 1). The activation of β-cell autoreactive T cells occurs in the pancreatic lymph node (PLN) due to autoantigen presentation by dendritic cells that migrated from the pancreas (Katsarou et al. 2017). As such, we also analyzed immune cell populations in the PLN of WT and miR146a-KO NOD mice and found no significant differences in CD4^+^ and CD8^+^ lymphocytes and CD11b^+^ and CD11c^+^ cells (Fig. 3*C*, Supplemental Fig. 1). Remarkedly, despite no observed differences in the immune cell populations in the thymus, spleen, and PLN, we observed a significant decrease in both CD4^+^ and CD8^+^ lymphocytes in the pancreas of miR146a-KO NOD mice (Fig. 3*D*). No differences were observed in CD11b^+^ and CD11c^+^ cells in the pancreas of WT NOD and miR146a-KO NOD mice (Supplemental Fig. 1). Taken together, these results clearly demonstrate that, while miRNA146 deletion has no major consequences on the frequency of immune cell populations in the thymus, spleen, and PLN in NOD mice, miR146a deletion results in a significant reduction in T lymphocytes in the pancreas of NOD mice.

**Figure 3.**
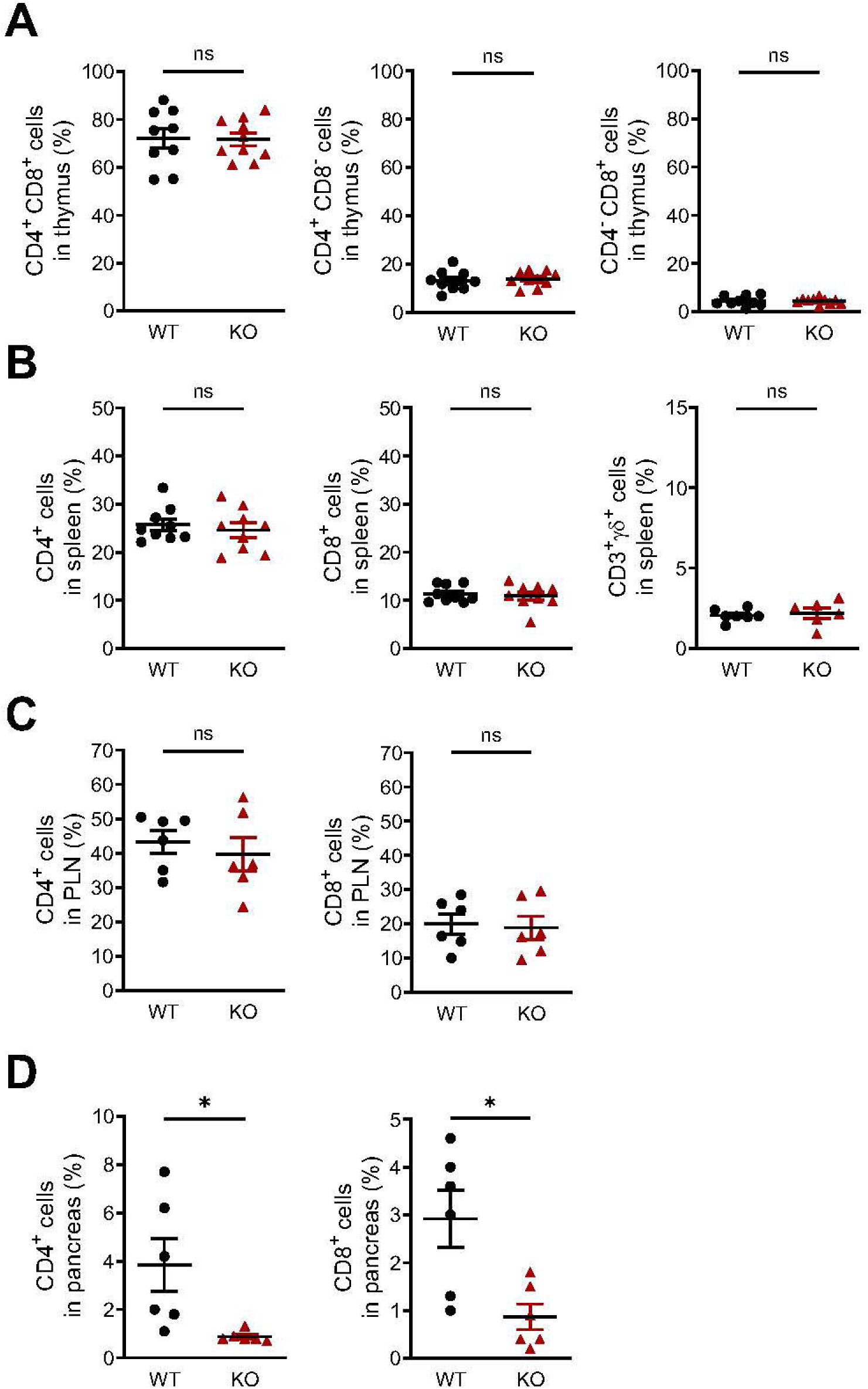
MicroRNA146a knockout results in significantly decreased T cells in the pancreas of NOD mice. Total cells were isolated from the thymus (A), spleen (B), PLN (C), and pancreas (D) of wildtype (WT) NOD and miR146a-knockout (KO) NOD mice and analyzed with flow cytometry for the various cell populations. Unpaired t-test, ns = not significant, * p < 0.05. Data is represented as mean ± SEM for n=6-9.

### MicroRNA146a knockout NOD mice are protected from spontaneous autoimmune diabetes and insulitis development

Next, we studied diabetes development in wildtype and miR146a-KO NOD female mice by monitoring blood glucose levels every week from 8-35 weeks of age and biweekly when the glucose level reached 250 mg/dL. Mice showing three consecutive readings of greater than 250 mg/dL were considered diabetic. Consistent with known literature for female NOD mice (Chen et al. 2020), 65% of NOD WT mice in our colony developed spontaneous diabetes by the age of 12-16 weeks. In contrast, only 20% of miR146a-KO NOD mice developed spontaneous autoimmune diabetes by 12-16 weeks of age. By the 35-week timepoint, only 40% of miR146a-KO NOD mice developed autoimmune diabetes, compared to over 80% of NOD WT mice (Fig. 4*A*). Histological analysis of the pancreas showed a significant decrease in immune cell infiltration into the pancreatic islets of miR146a-KO NOD mice (Fig. 4*B*). Overall insulitis score (Fig. 4*C*) and the percentage of infiltrated islets (Fig. 4*D*) were also significantly reduced in the absence of miR146a. In 8-week-old WT NOD mice, approximately 70% of islets were in stages 1-3 of insulitis, while in age-matched miR146a-KO NOD mice only about 35% of islets showed insulitis and about 65% of islets were in stage 0 of insulitis (Fig. 4*D*). Altogether, these data indicate that absence of miR146a protects NOD mice from hyperglycemia, insulitis, and autoimmune diabetes.

**Figure 4.**
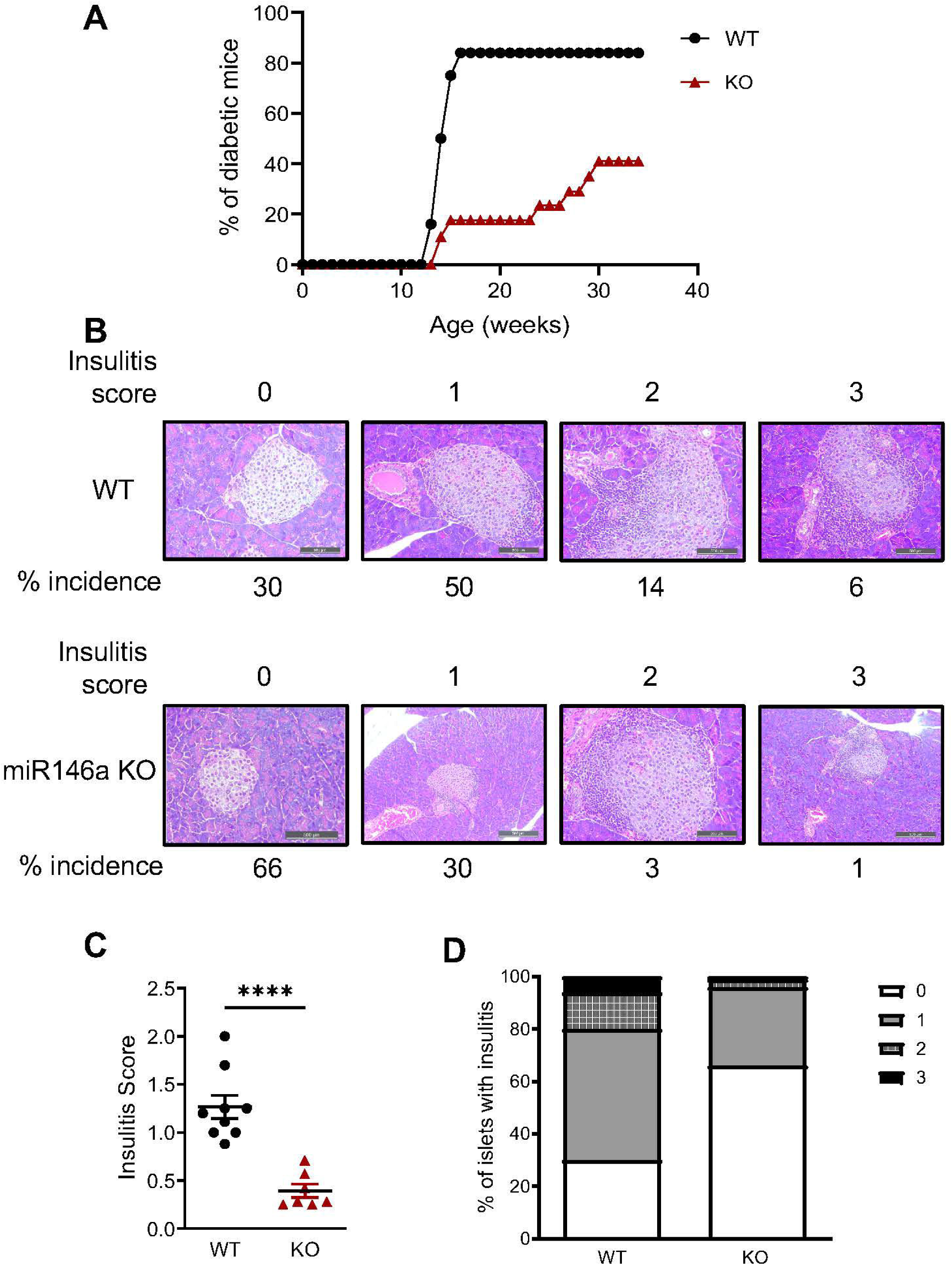
Spontaneous autoimmune diabetes and insulitis are reduced in the absence of miR146a in NOD mice. (A). Blood glucose was monitored weekly and then biweekly when levels reached 250 mg/dL. Mice with three consecutive readings of greater than 250 mg/dL were declared diabetic. n=26 (WT) and 28 (KO). (B). Pancreases from 8-week-old wildtype (WT) NOD and miR146a knockout (KO) NOD mice were analyzed by H&E staining for insulitis. Scale bar is 500 µM. (C). Mean insulitis score for pancreas of 8-week-old WT and KO mice. Unpaired t-test, **** p<0.0001 (D). Percentage of pancreatic islets at each stage of insulitis. No infiltration = 0 (white); Peri-islet infiltration of <25% of islet = 1 (gray); infiltration of 25-50% of islet = 2 (dashed); infiltration >50% of islet = 3 (black). (C). Data is represented as mean ± SEM for n=7-9.

### Absence of miR146a increases T regulatory cell number and function by increasing c-Rel expression and c-Rel binding to the FOXP3 promoter

Treg cells play a critical role in suppressing autoreactive T cells and maintaining self-tolerance in T1D (Hull, Peakman, and Tree 2017). To understand how absence of miR146a affects Treg cells in NOD mice, we examined the CD4^+^ FOXP3^+^ Treg cell populations in the thymus, spleen, and pancreatic lymph node (PLN). We found significantly increased T regulatory cells in the spleen and PLN of miR146a-KO NOD mice compared to WT NOD mice (Fig. 5*A*). We next examined how the absence of miR146a affects Treg cell-mediated suppression of T-cell receptor (TCR) activation-dependent CD4^+^ T cell proliferation. WT and miR146a-KO CD4^+^CD25^-^ T cells cultured alone showed no significant differences in proliferation. We observed miR146a-KO Treg cells exhibited greater suppressor function on the proliferation of both WT and miR146a-KO CD4^+^CD25^-^ T cells compared to WT Treg cells (Fig. 5*B-C*, Supplemental Fig. 2). These results suggest deletion of miR146a in NOD mice increases Treg cell number and suppressor function.

**Figure 5.**
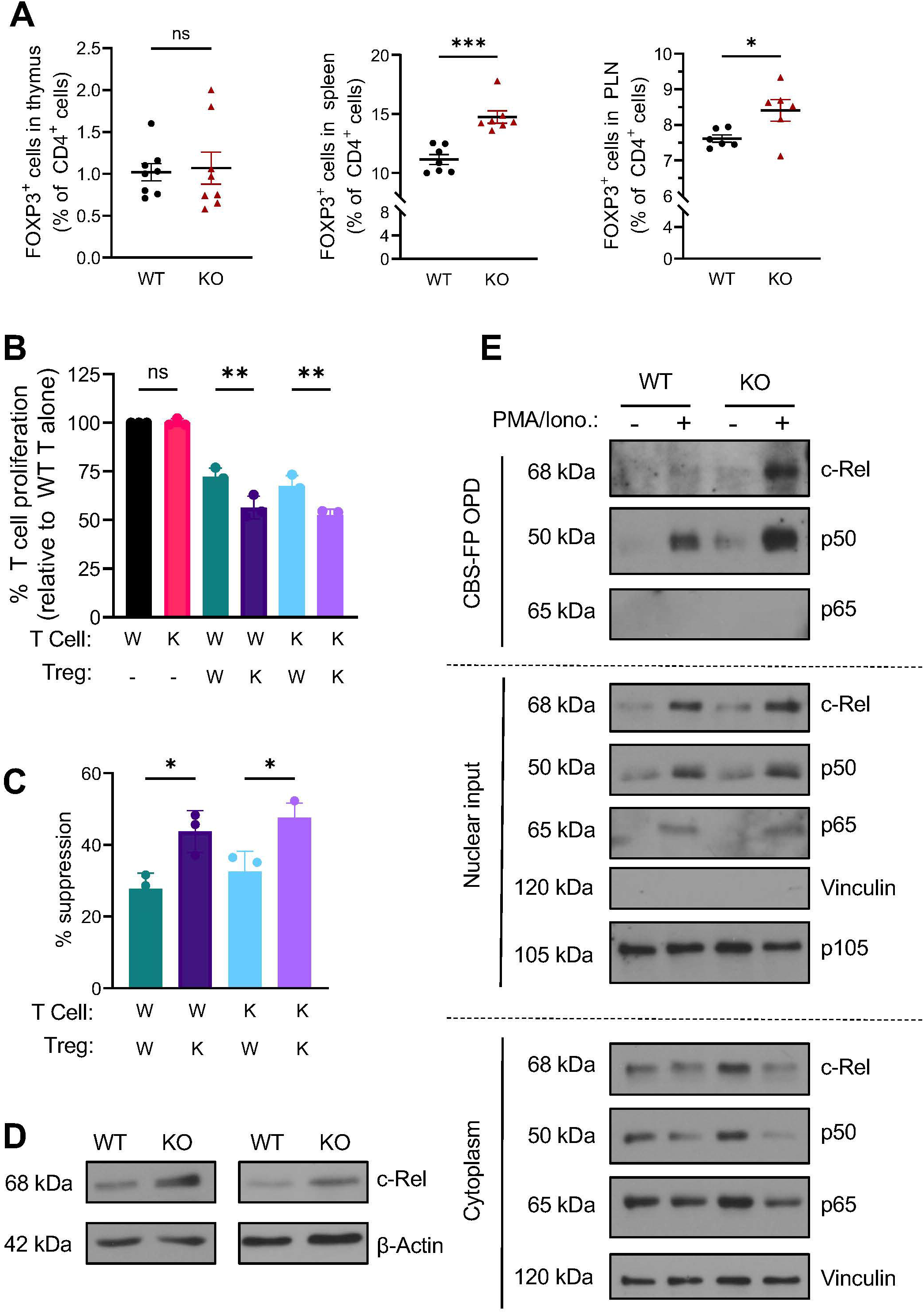
Absence of miR146a results in increased T regulatory cells that have enhanced suppressor function on T cell proliferation by modulating c-Rel. (A). Total cells were isolated from the thymus, spleen, and pancreatic lymph node of wildtype (WT) NOD and miR146a knockout (KO) NOD mice and analyzed with flow cytometry for FOXP3-positive T regulatory cells. Unpaired t-test, *p<0.05, ***p<0.001. Data is represented as mean ± SEM for n=3-7. (B and C). WT or miR146a KO CD4^+^ T cells were stained with CFSE and cultured with or without WT or miR146a-KO T regulatory cells for 72 hours with anti-CD3 and anti-CD28 stimulation. CFSE dye dilution was determined using flow cytometry. T cell proliferation was normalized to WT T cells alone over three independent experiments (B). Percent suppression was calculated as (% proliferation T cell alone - % proliferation T cell + Treg) / % proliferation T cell alone (C). (B and C). One-way ANOVA with multiple comparisons, *P<0.05, **P<0.01, ns = not significant. (D). CD4^+^CD25^+^ T regulatory cells from the spleen of WT or miR146a-KO mice were lysed and examined by Western blotting using antibodies against c-Rel. Actin was used as a loading control. Data is representative of n=5. (E) Cytoplasmic and nuclear lysate was obtained from CD4^+^CD25^+^ T regulatory cells from the spleen of WT or miR146a-KO mice, and nuclear lysate was used in an oligonucleotide pulldown assay (OPD) using biotinylated oligonucleotides corresponding to the c-Rel binding site in the mouse FOXP3 promoter (CBS-FP) oligonucleotide. Western blot was done on OPD sample, nuclear input, and cytoplasmic input against indicated antibodies. Data is representative of n=3.

NF-κB c-Rel is the major transcription factor that controls Treg cell development and peripheral function (Ruan et al. 2009; Visekruna et al. 2010; Fulford et al. 2021). Additionally, miR146a has previously been demonstrated to directly target c-Rel mRNA in B cells (Cho et al. 2018). We examined if c-Rel expression is altered in the absence of miR146a and found a robust increase in basal c-Rel protein level in CD4^+^CD25^+^ Treg cells isolated from the spleen of miR146a-KO NOD mice compared to WT NOD mice (Fig. 5*D*). NF-κB c-Rel binds to the FOXP3 promoter and orchestrates the formation of a complex that drives FOXP3 expression, thereby regulating Treg cell development (Ruan et al. 2009; De Jesus et al. 2021; Long et al. 2009). We examined whether the increase in c-Rel in miR146a-KO Treg cells results in its increased binding to the FOXP3 promoter by an oligonucleotide pulldown assay (OPD) using biotinylated oligonucleotides corresponding to the c-Rel binding site in the mouse FOXP3 promoter (CBS-FP) (De Jesus et al. 2021). We found that nuclear extracts of miR146a-KO Treg cells showed robustly increased TCR-activation dependent c-Rel binding to the FOXP3 promoter sequence compared to WT Treg cells (Fig. 5*E*). The binding of p50 subunit to the FOXP3 promoter was also increased in miR146a-KO Treg cells compared to WT Treg cells, suggesting potential c-Rel/p50 dimer containing transcriptional complex. NF-κB p65 binding was not detectable at this FOXP3 promoter region under our experimental conditions, which is in line with our previous study (De Jesus et al. 2021). The differences in c-Rel and p50 binding to the FOXP3 promoter are not simply due to increased protein levels as c-Rel and p50 levels were not drastically different in the nuclear fraction of activated WT and miR146a-KO Treg cells (Fig. 5*E*). In accordance with data in Fig 5D, basal levels of c-Rel were increased in the cytoplasmic fraction of miR146a-KO Treg cells. These data suggest a specific enrichment of active c-Rel that binds to the FOXP3 promoter in miR146a-KO Treg cells. Overall, our findings suggest that absence of miR146a in Treg cells results in increased c-Rel expression, and its binding at the FOXP3 promoter, which promotes Treg cell development and function in miR146a-knockout NOD mice.

## Discussion

In this study, we discovered the role of miR146a in regulating c-Rel-dependent Treg cell development and function relevant to autoimmune diabetogenesis in NOD mice. In the absence of miR146a, c-Rel expression and c-Rel binding to the FOXP3 promoter is enhanced. Increased c-Rel binding to the FOXP3 promoter contributes to increased Treg cell number and suppressor function, resulting in maintenance of self-tolerance and prevention of autoimmunity in miR146a-KO NOD mice. Our findings suggest increased miR146a expression in T1D patients (Fig. 1*A*) may contribute to autoimmune diabetes through compromising c-Rel-dependent FOXP3 expression, which controls Treg cell development and function. Under homeostatic conditions, a reciprocal relationship exists between miR146a and c-Rel with NF-κB c-Rel promoting miR146a expression (Yang et al. 2012), and miR146a serving as a feedback regulator of c-Rel protein levels ((Cho et al. 2018), Fig. 5*D*). This reciprocal relationship allows for maintenance of homeostatic immune cell function. Our study suggests that, in patients with T1D, hyperglycemia and enhanced O-GlcNAcylation may upregulate miR146a expression in T cells (Fig. 1*B-F*), potentially disrupting the homeostatic regulation of c-Rel by miR146a and contributing to a loss of self-tolerance seen in patients with T1D. Our study using miR146a-KO NOD mice shows that blocking miR146a may prevent loss of self-tolerance by increasing c-Rel-dependent FOXP3 expression, and thereby promoting Treg cell mediated maintenance of self-tolerance and preventing autoimmunity in T1D.

Our findings align with several studies suggesting that miR146a plays a pathogenic role in autoimmune conditions. Overexpression of miR146a in a transgenic mouse model results in a phenotype similar to autoimmune lymphoproliferative syndrome and in accumulation of germinal center B cells due to downregulation of Fas (Guo et al. 2013). In a mouse model of heart transplant rejection, conditional knockout of miR146a in Treg cells is protective, leading to increased allograft survival time and decreased T cell infiltration into the transplant (Jian Lu et al. 2021). Silencing miRNA146a in B cells ameliorated experimental myasthenia gravis (Zhang et al. 2015). In this study, we discovered miR146a-KO NOD mice are significantly protected from T1D (Fig. 3-4). Altogether, these studies indicate a potential role of miR146a in promoting autoimmunity and the utility of targeting miR146a as a therapeutic option in autoimmune conditions.

Furthermore, miR146a levels are increased in multiple autoimmune conditions in humans, including rheumatoid arthritis (Nakasa et al. 2008), psoriasis (Sonkoly et al. 2007), and myasthenia gravis (Jiayin Lu et al. 2013). microRNA146a is also elevated in the serum of T1D patients compared to age- and sex-matched healthy controls (Garavelli et al. 2020) and in NOD mice as they age and develop insulitis (Roggli et al. 2010). In this study, we discovered significantly increased miR146a expression in PBMCs of T1D patients compared to healthy controls (Fig. 1*A*). Thus, it appears that increased miR146a expression is a potential marker of T1D, which warrants further validation through large scale clinical studies.

While our study and others have observed a significant increase in miR146a expression in T1D patients in serum (Garavelli et al. 2020) and PBMCs (Fig. 1*A*), the molecular mechanisms responsible for the upregulation were unclear. We identified both hyperglycemia and O-GlcNAcylation as physiological regulators of miR146a expression in T cells (Fig. *1B*, *1F*). Thus, we propose hyperglycemia and enhanced cellular O-GlcNAcylation are mechanistic explanations for the increased miR146a observed in T1D patients. It remains to be addressed which transcription factor(s) are responsible for the O-GlcNAc-dependent changes in miR146a expression. The promotor of miR146a contains putative binding sites for NF-κB, IRF3/7, and C/EBPβ, but LPS-induced miR146a expression was only dependent NF-κB in myeloid cells (Taganov et al. 2006). In T cells, both basal and TCR-induced miR146a expression is dependent on c-Rel and p50. The contribution of p65 was not studied (Yang et al. 2012). NF-κB p65 has multiple sites of O-GlcNAcylation, while NF-κB c-Rel has one site of O-GlcNAcylation at serine 350 (Liu and Ramakrishnan 2021). Future studies should explore how blocking O-GlcNAcylation at specific residues on proteins affects miR146a expression.

Previously, miR146a-KO C57BL/6 × 129/sv mice were observed to develop myeloproliferation with age around 14 months old (Boldin et al. 2011). We did not observe any changes in major hematological parameters and immune cell populations in 2–3-month-old miR146a-KO NOD mice (Fig. 2-3). Future studies are needed to explore whether miR146a-KO NOD mice also develop myeloproliferation with age, or if there are potentially strain-specific differences.

Since the discovery of microRNAs in 1993, many microRNAs have been implicated in disease development (Bhaskaran and Mohan 2013). In regard to autoimmune diabetes, microRNAs play a role in pancreatic β-cell function, insulin synthesis, and immune system homeostasis (Margaritis et al. 2021). It is possible that protection from autoimmune diabetes in miR146a-KO NOD mice may also result from the function of miR146a in different hematopoietic cells and in pancreatic β-cells. In the Min6 pancreatic β-cell line, inhibition of miR146a reduced cytokine-induced β-cell death, while overexpression of miR146a increased β-cell death (Roggli et al. 2010). The mechanisms by which miR146a may regulate β-cell death and the relevance of this phenotype *in vivo* is unclear. For example, it remains to be explored whether miR146a inhibition could protect pancreatic β-cells from cell death mediated by autoreactive CD8^+^ T cells versus cytokine mediated cell death. In miR146a-KO NOD mice, the marked reduction in T cell infiltration into the pancreas suggests the mechanism for diabetes protection in miR146a-KO NOD mice proceeds T cell infiltration into the pancreas and as such, likely excludes a mechanism solely intrinsic to β-cell apoptosis. To overcome some of the limitations with global miR146a-KO NOD mice, we evaluated the specific function of miR146a in Treg cells using *ex vivo* assays. In the coculture suppression assay, the enhanced suppressor function of miR146a-KO Treg cells was observed regardless of the WT or miR146a-KO CD4^+^CD25^-^ cell genotype (Fig. 5*B-C*), suggesting our findings are intrinsic to the Treg cells. However, we acknowledge the limitations of the current global miR146a-KO NOD mouse model, and future studies using Treg cell-specific deletion of miR146a in NOD mice are necessary to fully understand the contribution of miR146a in Treg cells to protection from autoimmune diabetes.

In addition to IRAK1 and TRAF6, NF-κB c-Rel was identified as a direct target of miR146a in B cells (Cho et al. 2018). In T regulatory cells, c-Rel binds to the FOXP3 promoter and orchestrates the formation of a transcription factor complex that drives *FOXP3* expression (Ruan et al. 2009; Long et al. 2009). T regulatory cell induction is significantly impaired in the absence of c-Rel (Visekruna et al. 2010; Ruan et al. 2009), and overexpression of c-Rel significantly increases FOXP3 promoter activity (Ruan et al. 2009). Furthermore, c-Rel promotes the peripheral suppressor function of T regulatory cells with absence of c-Rel resulting in diminished frequency of Treg cells that express CD103, ICOS, and TIGIT (Fulford et al. 2021). miR146a-KO Treg cells showed a robust increase in c-Rel and its binding to the FOXP3 promoter, which correlated with increased Treg cells in miR146a-KO NOD mice (Fig. 5). These data mechanistically suggest a miR146a-c-Rel axis contributes to Treg cell development and function. While unstimulated miR146a-KO Treg cells had increased c-Rel protein levels in the cytoplasm, the levels of translocated c-Rel in the nucleus of stimulated cells were similar between WT and miR146a-KO Treg cells. This finding suggests increased c-Rel binding to the FOXP3 promoter in miR146a-KO Treg cells is likely not due to simply increased c-Rel protein levels in the nucleus, but rather absence of miR146a in Treg cells poises a subcellular fraction of c-Rel for increased binding to the FOXP3 promoter. Further studies are needed to delineate how c-Rel binding to the FOXP3 promoter is enhanced without a significant increase in total nuclear c-Rel level and whether it binds as a homodimer or a heterodimer with p50 to enhance FOXP3 transcription. Irrespective of such molecular details, this study further strengthens the importance of c-Rel in T regulatory cell development and function and suggests targeting miR146a may be a strategy to increase c-Rel-dependent Treg function to restore self-tolerance in autoimmune conditions.

Restoring self-tolerance addresses an unmet need as the incidence of T1D is constantly rising globally, with an annual increase of 3-4% (Akil et al. 2021), with no cure. Insulin replacement and beta cell replacement therapies do not address the underlying autoimmunity present in T1D. Defective Treg cell function is identified even in early stages of T1D before clinical symptoms are observed, suggesting that Treg cell dysfunction plays a causative role in T1D pathogenesis, rather than being an effect of T1D (Hull, Peakman, and Tree 2017). Several clinical therapies aimed at increasing Treg cell number and function, including low-dose IL2 administration and adoptive Treg cell therapy have not yet yielded expected success rates likely due to multiple factors (Hull, Peakman, and Tree 2017). For example, Treg cells are generally isolated from peripheral blood and then expanded *ex vivo* for supplementation therapy, which raises concerns of Treg cells losing *FOXP3* expression and lineage stability or Treg cells obtaining an exhausted, non-suppressive phenotype overtime (Bettini and Bettini 2021). Additionally, multiple studies have indicated that Treg cells obtain functional specialization depending on the tissue site, and as such, it is uncertain if peripheral blood Treg cells will have optimal localization to and suppressor function in the pancreatic lymph node and pancreas (Bettini and Bettini 2021; Hull, Peakman, and Tree 2017). Our findings suggest that miR146a inhibition may be a viable therapeutic strategy to overcome the current challenges with Treg therapies as miR146a inhibition may specifically increase the frequency of Treg cells in the pancreatic lymph node in the context of T1D and enhance the suppressor function of Treg cells.

## Contributions

CNA, SB, ARL, and PR conceived and planned the experiments. CNA, SB, ARL, KN, and PR performed experiments and interpretated the results. CNA, SB, and PR wrote the original manuscript. All authors reviewed the final manuscript.

## Acknowledgments

We thank Dr. Mark Boldin and Dr. Jimmy Zhao for valuable discussions on miR-146a biology. We are grateful to past lab members Dr. Joshua T Centore, Dr. Li Liu, and Ms. Julia Hluck for their support in genotyping mice and discussions.

## Funding

PR was supported by Breakthrough T1D (formerly JDRF) grants 3-SRA-2022–1193-S-B and 3-SRA-2024–1552-S-B, NIH/NIAID grants R01AI116730 and R21AI144264, NIH/NIDDK R01DK128463, and VA BLRD grant I01 BX005941. CNA and ARL were supported NIH/NEI T32 predoctoral training grant EY007157 through the Department of Ophthalmology by the NIH MSTP T32 GM152319. CNA was also supported by the NIH/NIDDK F30 Kirschstein-NRSA predoctoral fellowship grant F30DK143695.

## Supplemental Figure Legends

**Supplemental Figure 1. MicroRNA146a knockout does not affect myeloid and B cell populations in NOD mice.** Total cells were isolated from the spleen (A), PLN (B), and pancreas (C) of wildtype (WT) NOD and miR146a-knockout (KO) NOD mice and analyzed with flow cytometry for the various cell populations. Unpaired t-test, ns = not significant. Data is represented as mean ± SEM for n=4-8.

**Supplemental Figure 2. Absence of miR146a results in T regulatory cells that have enhanced suppressor function on T cell proliferation.** WT or miR146a KO CD4^+^ T cells were stained with CFSE and cultured with or without WT or miR146a-KO T regulatory cells for 72 hours with anti-CD3 and anti-CD28 stimulation. CFSE dye dilution was determined using flow cytometry. Representative flow cytometry plots are shown. Data is representative of n=3.

## Notes

### Competing Interest Statement

The authors have declared no competing interest.

## References

Akil, Ammira Al Shabeeb, Esraa Yassin, Aljazi Al-Maraghi, Elbay Aliyev, Khulod Al-Malki, and Khalid A. Fakhro. 2021. “Diagnosis and Treatment of Type 1 Diabetes at the Dawn of the Personalized Medicine Era.” Journal of Translational Medicine 19 (1): 137. 10.1186/s12967-021-02778-6.

Assmann, Taís S., Guilherme C.K. Duarte, Letícia A. Brondani, Pedro H.O. de Freitas, Égina M. Martins, Luís H. Canani, and Daisy Crispim. 2017. “Polymorphisms in Genes Encoding MiR-155 and MiR-146a Are Associated with Protection to Type 1 Diabetes Mellitus.” Acta Diabetologica 54: 433–41. 10.1007/s00592-016-0961-y.

Bettini, Maria, and Matthew L. Bettini. 2021. “Function, Failure, and the Future Potential of Tregs in Type 1 Diabetes.” Diabetes 70 (6): 1211–19. 10.2337/DBI18-0058.

Bhaskaran, M, and M Mohan. 2013. “MicroRNAs: History, Biogenesis, and Their Evolving Role in Animal Development and Disease.” Vet Pathol 51 (4): 759–74. 10.1177/0300985813502820.

Boldin, Mark P., Konstantin D. Taganov, Dinesh S. Rao, Lili Yang, Jimmy L. Zhao, Manorama Kalwani, Yvette Garcia-Flores, et al. 2011. “MiR-146a Is a Significant Brake on Autoimmunity, Myeloproliferation, and Cancer in Mice.” Journal of Experimental Medicine 208 (6): 1189–1201. 10.1084/jem.20101823.

Chen, Dawei, Terri C Thayer, Li Wen, and F Susan Wong. 2020. “Mouse Models of Autoimmune Diabetes: The Nonobese Diabetic (NOD) Mouse.” Methods in Molecular Biology 2128: 87–92. 10.1007/978-1-0716-0385-7_6.

Cho, Sunglim, Hyang Mi Lee, I. Shing Yu, Youn Soo Choi, Hsi Yuan Huang, Somaye Sadat Hashemifar, Ling Li Lin, et al. 2018. “Differential Cell-Intrinsic Regulations of Germinal Center B and T Cells by MiR-146a and MiR-146b.” Nature Communications 9 (2757). 10.1038/s41467-018-05196-3.

Dennis, Michael D., Tabitha L. Schrufer, Sarah K. Bronson, Scot R. Kimball, and Leonard S. Jefferson. 2011. “Hyperglycemia-Induced O-GlcNAcylation and Truncation of 4E-BP1 Protein in Liver of a Mouse Model of Type 1 Diabetes.” Journal of Biological Chemistry 286 (39): 34286–97. 10.1074/jbc.M111.259457.

Fulford, Thomas S., Raelene Grumont, Rushika C. Wirasinha, Darcy Ellis, Adele Barugahare, Stephen J. Turner, Haroon Naeem, et al. 2021. “C-Rel Employs Multiple Mechanisms to Promote the Thymic Development and Peripheral Function of Regulatory T Cells in Mice.” European Journal of Immunology 51 (8): 2006–26. 10.1002/eji.202048900.

Garavelli, Silvia, Sara Bruzzaniti, Elena Tagliabue, Francesco Prattichizzo, Dario Di Silvestre, Francesco Perna, Lucia La Sala, et al. 2020. “Blood Co-Circulating Extracellular Micrornas and Immune Cell Subsets Associate with Type 1 Diabetes Severity.” International Journal of Molecular Sciences 21 (2): 477. 10.3390/ijms21020477.

Gebert, Luca F.R., and Ian J. MacRae. 2019. “Regulation of MicroRNA Function in Animals.” Nature Reviews Molecular Cell Biology 20: 21–37. 10.1038/s41580-018-0045-7.

Guo, Qiuye, Jinjun Zhang, Jingyi Li, Liyun Zou, Jinyu Zhang, Zunyi Xie, Xiaolan Fu, et al. 2013. “Forced MiR-146a Expression Causes Autoimmune Lymphoproliferative Syndrome in Mice via Downregulation of Fas in Germinal Center B Cells.” Blood 121 (24): 4875–83. 10.1182/blood-2012-08-452425.

Hart, Gerald W. 2019. “Nutrient Regulation of Signaling and Transcription.” Journal of Biological Chemistry 294 (7): 2211–31. 10.1074/jbc.AW119.003226.

Hull, Caroline M., Mark Peakman, and Timothy I.M. Tree. 2017. “Regulatory T Cell Dysfunction in Type 1 Diabetes: What’s Broken and How Can We Fix It?” Diabetologia 60 (10): 1839–50. 10.1007/s00125-017-4377-1.

Iacona, Joseph R., and Carol S. Lutz. 2019. “MiR-146a-5p: Expression, Regulation, and Functions in Cancer.” Wiley Interdisciplinary Reviews: RNA 10 (4): e1533. 10.1002/wrna.1533.

Jesus, Tristan J. De, Jeffrey A. Tomalka, Joshua T. Centore, Franklin D. Staback Rodriguez, Ruchira A. Agarwal, Angela R. Liu, Timothy S. Kern, and Parameswaran Ramakrishnan. 2021. “Negative Regulation of FOXP3 Expression by C-Rel O-GlcNAcylation.” Glycobiology 31 (7): 812–26. 10.1093/glycob/cwab001.

Katsarou, Anastasia, Soffia Gudbjörnsdottir, Araz Rawshani, Dana Dabelea, Ezio Bonifacio, Barbara J. Anderson, Laura M. Jacobsen, Desmond A. Schatz, and Ake Lernmark. 2017. “Type 1 Diabetes Mellitus.” Nature Reviews Disease Primers 3: 17016. 10.1038/nrdp.2017.16.

Kazemi, Zahra, Hana Chang, Sarah Haserodt, Cathrine McKen, and Natasha E. Zachara. 2010. “O-Linked β-N-Acetylglucosamine (O-GlcNAc) Regulates Stress-Induced Heat Shock Protein Expression in a GSK-3β-Dependent Manner.” Journal of Biological Chemistry 285 (50): 39096–107. 10.1074/jbc.M110.131102.

Liu, Angela Rose, and Parameswaran Ramakrishnan. 2021. “Regulation of Nuclear Factor-KappaB Function by O-GlcNAcylation in Inflammation and Cancer.” Frontiers in Cell and Developmental Biology 9 (751761). 10.3389/fcell.2021.751761.

Long, Meixiao, Sung Gyoo Park, Ian Strickland, Matthew S. Hayden, and Sankar Ghosh. 2009. “Nuclear Factor-ΚB Modulates Regulatory T Cell Development by Directly Regulating Expression of Foxp3 Transcription Factor.” Immunity 31 (6): 921–31. 10.1016/j.immuni.2009.09.022.

Lu, Jian, Weiwei Wang, Peiyuan Li, Xiaodong Wang, Chao Gao, Baotong Zhang, Xuezhi Du, Yanhong Liu, Yong Yang, and Feng Qi. 2021. “MiR-146a Regulates Regulatory T Cells to Suppress Heart Transplant Rejection in Mice.” Cell Death Discovery 7: 165. 10.1038/s41420-021-00534-9.

Lu, Jiayin, Mei Yan, Yuzhong Wang, Junmei Zhang, Huan Yang, Fa fa Tian, Wenbin Zhou, Ning Zhang, and Jing Li. 2013. “Altered Expression of MiR-146a in Myasthenia Gravis.” Neuroscience Letters 555: 85–90. 10.1016/j.neulet.2013.09.014.

Lu, Li Fan, Mark P. Boldin, Ashutosh Chaudhry, Ling Li Lin, Konstantin D. Taganov, Toshikatsu Hanada, Akihiko Yoshimura, David Baltimore, and Alexander Y. Rudensky. 2010. “Function of MiR-146a in Controlling Treg Cell-Mediated Regulation of Th1 Responses.” Cell 142 (6): 914–29. 10.1016/j.cell.2010.08.012.

Mannino, Michael P., and Gerald W. Hart. 2022. “The Beginner’s Guide to O-GlcNAc: From Nutrient Sensitive Pathway Regulation to Its Impact on the Immune System.” Frontiers in Immunology 13 (828648). 10.3389/fimmu.2022.828648.

Margaritis, Kosmas, Georgia Margioula-siarkou, Styliani Giza, Eleni P. Kotanidou, Vasiliki Regina Tsinopoulou, Athanasios Christoforidis, and Assimina Galli-tsinopoulou. 2021. “Micro-RNA Implications in Type-1 Diabetes Mellitus: A Review of Literature.” International Journal of Molecular Sciences 22 (22): 12165. 10.3390/ijms222212165.

Mortazavi-Jahromi, Seyed Shahabeddin, Mona Aslani, and Abbas Mirshafiey. 2020. “A Comprehensive Review on MiR-146a Molecular Mechanisms in a Wide Spectrum of Immune and Non-Immune Inflammatory Diseases.” Immunology Letters 227: 8–27. 10.1016/j.imlet.2020.07.008.

Nakasa, Tomoyuki, Shigeru Miyaki, Atsuko Okubo, Megumi Hashimoto, Keiichiro Nishida, Mitsuo Ochi, and Hiroshi Asahara. 2008. “Expression of MicroRNA-146 in Rheumatoid Arthritis Synovial Tissue.” Arthritis and Rheumatism 58 (5): 1284–92. 10.1002/art.23429.

Park, Sung Kyun, Xiaorong Zhou, Kathryn E. Pendleton, Olga V. Hunter, Jennifer J. Kohler, Kathryn A. O’Donnell, and Nicholas K. Conrad. 2017. “A Conserved Splicing Silencer Dynamically Regulates O-GlcNAc Transferase Intron Retention and O-GlcNAc Homeostasis.” Cell Reports 20 (5): 1088–99. 10.1016/j.celrep.2017.07.017.

Perides, George, Eric R. Weiss, Emily S. Michael, Johanna M. Laukkarinen, Jeremy S. Duffield, and Michael L. Steer. 2011. “TNF-α-Dependent Regulation of Acute Pancreatitis Severity by Ly-6Chi Monocytes in Mice.” Journal of Biological Chemistry 286 (15): 13327–35. 10.1074/jbc.M111.218388.

Qian, Kevin, Simeng Wang, Minnie Fu, Jinfeng Zhou, Jay Prakash Singh, Min Dian Li, Yunfan Yang, et al. 2018. “Transcriptional Regulation of O-GlcNAc Homeostasis Is Disrupted in Pancreatic Cancer.” Journal of Biological Chemistry 293 (36): 13989–0. 10.1074/jbc.RA118.004709.

Rajendeeran, Anandi, and Klaus Tenbrock. 2021. “Regulatory T Cell Function in Autoimmune Disease.” Journal of Translational Autoimmunity 4: 100130. 10.1016/j.jtauto.2021.100130.

Roggli, Elodie, Aurore Britan, Sonia Gattesco, Nathalie Lin-Marq, Amar Abderrahmani, Paolo Meda, and Romano Regazzi. 2010. “Involvement of MicroRNAs in the Cytotoxic Effects Exerted by Proinflammatory Cytokines on Pancreatic β-Cells.” Diabetes 59 (4): 978–86. 10.2337/db09-0881.

Ruan, Qingguo, Vasumathi Kameswaran, Yukiko Tone, Li Li, Hsiou Chi Liou, Mark I. Greene, Masahide Tone, and Youhai H. Chen. 2009. “Development of Foxp3+ Regulatory T Cells Is Driven by the C-Rel Enhanceosome.” Immunity 31 (6): 932–40. 10.1016/j.immuni.2009.10.006.

Sandor, A. M., J. Jacobelli, and R. S. Friedman. 2019. “Immune Cell Trafficking to the Islets during Type 1 Diabetes.” Clinical and Experimental Immunology 198 (3): 314–25. 10.1111/cei.13353.

Sonkoly, Enikö, Tianling Wei, Peter C.J. Janson, Annika Sääf, Lena Lundeberg, Maria Tengvall-Linder, Gunnar Norstedt, et al. 2007. “MicroRNAs: Novel Regulators Involved in the Pathogenesis of Psoriasis?” PLoS ONE 2 (7): e610. 10.1371/journal.pone.0000610.

Taganov, Konstantin D., Mark P. Boldin, Kuang Jung Chang, and David Baltimore. 2006. “NF-ΚB-Dependent Induction of MicroRNA MiR-146, an Inhibitor Targeted to Signaling Proteins of Innate Immune Responses.” Proceedings of the National Academy of Sciences of the United States of America 103 (33): 12481–86. 10.1073/pnas.0605298103.

Truett, Gary E., P. Heeger, R. L. Mynatt, A. A. Truett, J. A. Walker, and M. L. Warman. 2000. “Preparation of PCR-Quality Mouse Genomic Dna with Hot Sodium Hydroxide and Tris (HotSHOT).” BioTechniques 29: 52–54. 10.2144/00291bm09.

Turley, Shannon J., Je-Wook Lee, Nick Dutton-Swain, Diane Mathis, and Christophe Benoist. 2005. “Endocrine Self and Gut Non-Self Intersect in the Pancreatic Lymph Nodes.” Proceedings of the National Academy of Sciences of the United States of America 102 (49): 17729–33. 10.1073/pnas.0509006102.

Visekruna, Alexander, Magdalena Huber, Anne Hellhund, Evita Bothur, Katharina Reinhard, Nadine Bollig, Nicole Schmidt, Thorsten Joeris, Michael Lohoff, and Ulrich Steinhoff. 2010. “C-Rel Is Crucial for the Induction of Foxp3+ Regulatory CD4 + T Cells but Not TH17 Cells.” European Journal of Immunology 40 (3): 671–76. 10.1002/eji.200940260.

Yang, Lili, Mark P. Boldin, Yang Yu, Claret Siyuan Liu, Chee Kwee Ea, Parameswaran Ramakrishnan, Konstantin D. Taganov, Jimmy L. Zhao, and David Baltimore. 2012. “MiR-146a Controls the Resolution of T Cell Responses in Mice.” Journal of Experimental Medicine 209 (9): 1655–70. 10.1084/jem.20112218.

Zhang, Jun Mei, Ge Jia, Qun Liu, Jue Hu, Mei Yan, Bai Feng Yang, Huan Yang, Wen Bin Zhou, and Jing Li. 2015. “Silencing MiR-146a Influences B Cells and Ameliorates Experimental Autoimmune Myasthenia Gravis.” Immunology 144: 56–67. 10.1111/imm.12347.

