## Supplementary figures and images for "MicroRNA-146a Deficiency Protects NOD Mice from Autoimmune Diabetes by Enhancing c-Rel-Dependent Regulatory T Cell Function"

### Supplemental fig 1

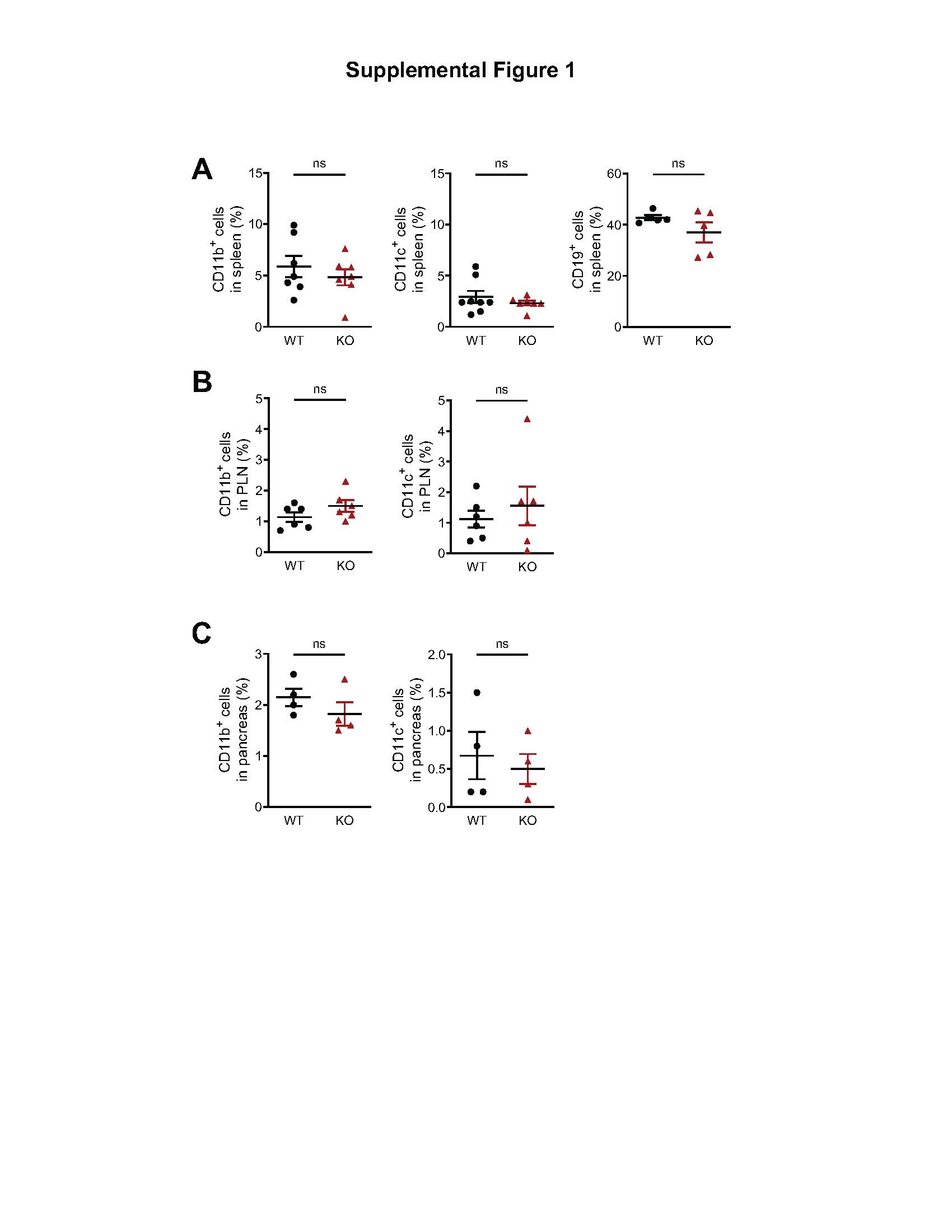

### Supplemental fig 2

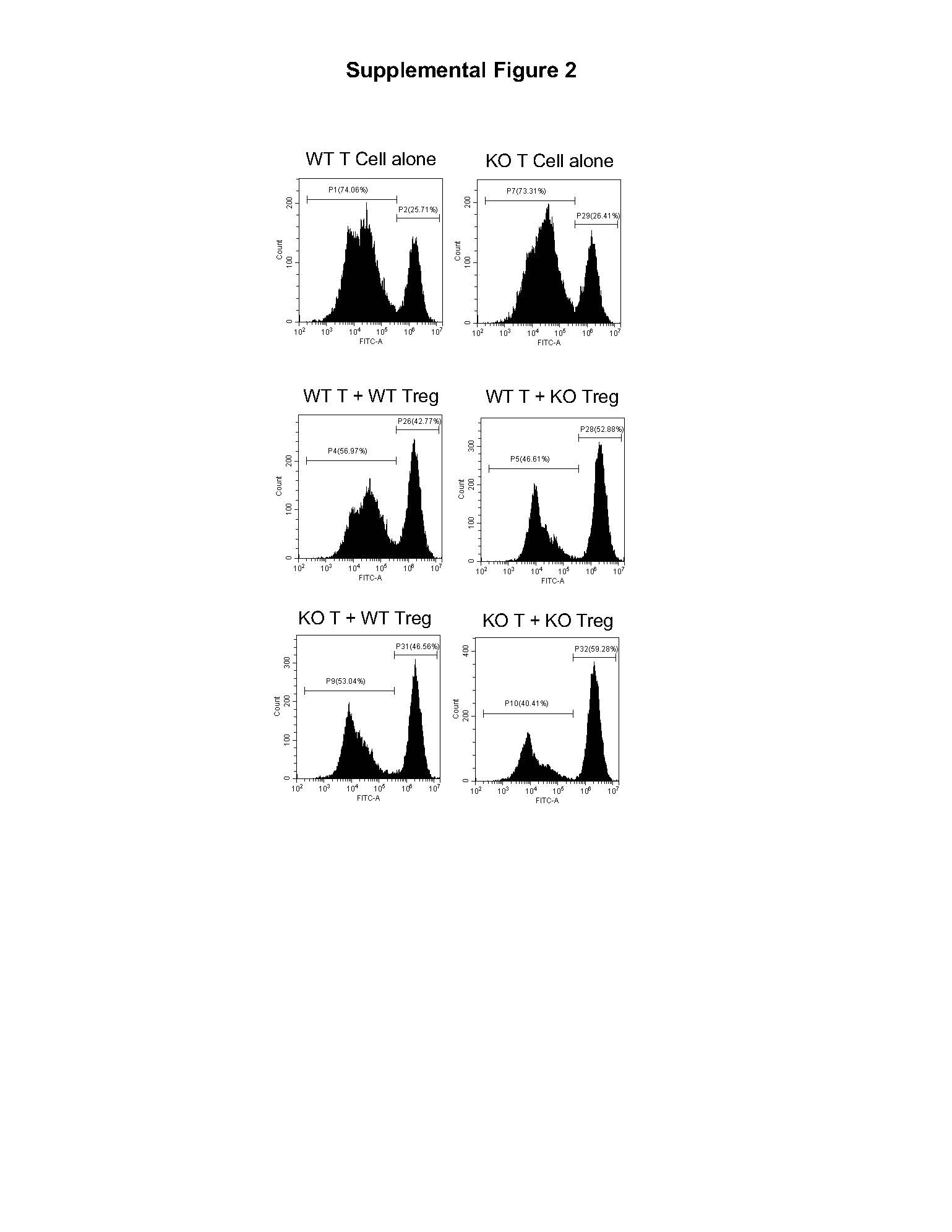
