## Supplemental table 1 for "MicroRNA-146a Deficiency Protects NOD Mice from Autoimmune Diabetes by Enhancing c-Rel-Dependent Regulatory T Cell Function"

| **Parameter** | **WT (n=4)** | **KO (n=4)** | **P value** |
| --- | --- | --- | --- |
| Lymphocyte (unit) | 4.13 ± 0.51 | 3.73 ± 1.85 | 0.72175781 |
| Monocyte (unit) | 0.60 ± 0.08 | 0.60 ± 0.20 | 1 |
| Granulocyte (unit) | 2.28 ± 0.48 | 1.85 ± 0.70 | 0.393641983 |
| RBC Time (unit) | 14.03 ± 0.05 | 14.03 ± 0.05 | 1 |
| HGB (g/dL) | 14.73 ± 0.37 | 15.23 ± 0.70 | 0.301667375 |
| HCT (unit) | 41.78 ± 1.16 | 43.43 ± 2.22 | 0.282390609 |
| MCV (unit) | 48.83 ± 0.24 | 48.48 ± 0.28 | 0.129828993 |
| RDWa (unit) | 34.78 ± 0.62 | 34.65 ± 0.58 | 0.79411804 |
| RDW % | 18.13 ± 0.30 | 18.28 ± 0.16 | 0.449284004 |
| MCHC (unit) | 35.23 ± 0.25 | 35.05 ± 0.44 | 0.555814775 |
| MCH (unit) | 17.20 ± 0.13 | 17.00 ± 0.26 | 0.268595288 |
| WBC Time (unit) | 9.18 ± 0.05 | 9.20 ± 0.00 | 0.391002219 |
| MPV (unit) | 6.23 ± 0.24 | 6.23 ± 0.19 | 1 |

**Supplemental Table 1. MicroRNA146a knockout does not lead to significant changes in peripheral blood parameters in NOD mice.** Blood samples from NOD wildtype (WT) and miR146a knockout (KO) were analyzed in the Hemavet blood analyzer. Unpaired t-test was used to calculate p-values. Data is represented as mean ± SEM for n=4.
